# SYBR Gold staining of single stranded nucleic acids is strongly influenced by the presence of guanosines

**DOI:** 10.64898/2026.05.19.722025

**Authors:** Jakob Rien, Mauro Zimmermann, Jonathan Hall, Mihaela Zavolan, Nitish Mittal

## Abstract

SYBR Gold is one of the most strongly fluorescent members of the SYBR family of asymmetrical cyanine dyes commonly used for staining double and single-stranded DNA and RNA. The dye fluorescence increases dramatically upon nucleic acid binding. The primary mode of binding to double-stranded DNA is through intercalation between the base-paired deoxyribonucleotides, while the precise mode of binding to single-stranded DNA and to RNA is less well understood. Here we show that the fluorescence activity of SYBR Gold is strongly influenced by the presence of guanines in single-stranded nucleic acids. We expect this insight to aid in the evaluation of SYBR Gold-based measurements of nucleic acid abundance and spark further studies into the binding behaviour of the SYBR family dyes.

## Introduction

Fluorescent nucleic acid-staining dyes - in particular from the SYBR family^1^ that includes SYBR Green I, SYBR Green II, and SYBR Gold - offer a straightforward way to visualize RNA and DNA and have become a routine reagent in a variety of assays. It is generally understood that SYBR Gold functions by intercalating between base-pairs of the DNA helix^1^, leading to a large increase in its fluorescent signal^1^. The first structural evidence for this interaction appeared in 2021, when the crystal structure of SYBR Gold was described, together with evidence for DNA intercalation^2^. However, a definitive binding mechanism for SYBR Gold remains to be determined, particularly regarding its interaction with single-stranded DNA and with RNA. During experiments in which we were studying guanosine oxidation in strands of RNA, we observed that SYBR Gold staining was surprisingly ineffective for the visualization of specific RNA oligonucleotides that could be detected by alternative methods. We determined that the key difference between RNAs that could and those that could not be visualized was their sequence composition: RNAs that lacked (unmodified) guanosines produced very weak fluorescence signals. These observations seemed at odds with the original published findings of Tuma et al.^1^, namely that after poly-rU, which yielded the highest intensity SYBR Gold signal, poly-rG and single-stranded poly-T DNA bind equally well. To resolve this discrepancy, we performed a series of experiments aimed at clarifying the effect to guanosine in mixed-sequence RNAs.

## Materials and Methods

### Oligonucleotides

Oligonucleotides were either synthesized in house or purchased from Microsyth AG Switzerland.

### PAGE gel

We prepared denaturing 14% urea PAGE gels by combining 28.8 g of urea, 21 mL of 40% acrylamide, 6 ml 10xTBE buffer (Tris/Borate/EDTA) and MiliQ water, added to a final volume of 60 mL. For each sample we used 500 ng of oligonucleotide in a final volume of 20 µL, including 2X RNA loading dye (95 % formamide, 0.025 % SDS, 0.025 % bromophenol blue and 1mM EDTA) and heated at 80° C for 5 min before loading onto the gel apparatus. The gel was run at 200-250V for 2.0–2.5 h, then incubated in 1xTBE buffer with 1:10’000 diluted SYBR Gold (ThermoFisher cat. S11494) for 10 min in the dark with constant shaking at low speed. Finally, the SYBR Gold fluorescence was visualized on an epiblue transilluminator (Dark Reader DR-88X by Clare Chemical Research). For non-denaturing or native PAGE gel, we omitted urea from the gel solution recipe mentioned above, used 6x loading dye (New England Biolabs# B70245) and did not heat samples before loading.

### dsDNA Generation

dsDNAs were generated by hybridizing complementary DNA strands. For each hybridization, we used 100 µM of each strand of the oligonucleotide. The hybridization occurred during cooling from 95°C to room temperature (approx. 2 h). We used a 2x DNA hybridization buffer (2 mM EDTA, 100 mM NaCl, 20 mM Tris pH7.5). The final concentration of the buffer after mixing with the samples was 1x. For dsDNA, we use non-denaturing PAGE gels.

### Image Analysis

The band intensities from SYBR Gold stains were analysed using ImageJ. The data were normalized to the strongest signal.

## Results

Our motivation for investigating the influence of guanosine (G) on SYBR Gold signal intensity came from an initial observation that iso-sequential RNA oligonucleotides containing predominantly oxidized guanosines (8-oxo-G) yielded significantly lower fluorescence signal intensities compared to those containing equivalent amounts of unmodified guanosine (compare lanes 9/10 vs. lanes 5/6; lanes 7/8 vs. lanes 3/4; **Fig. 1A**).

**Fig. 1.**
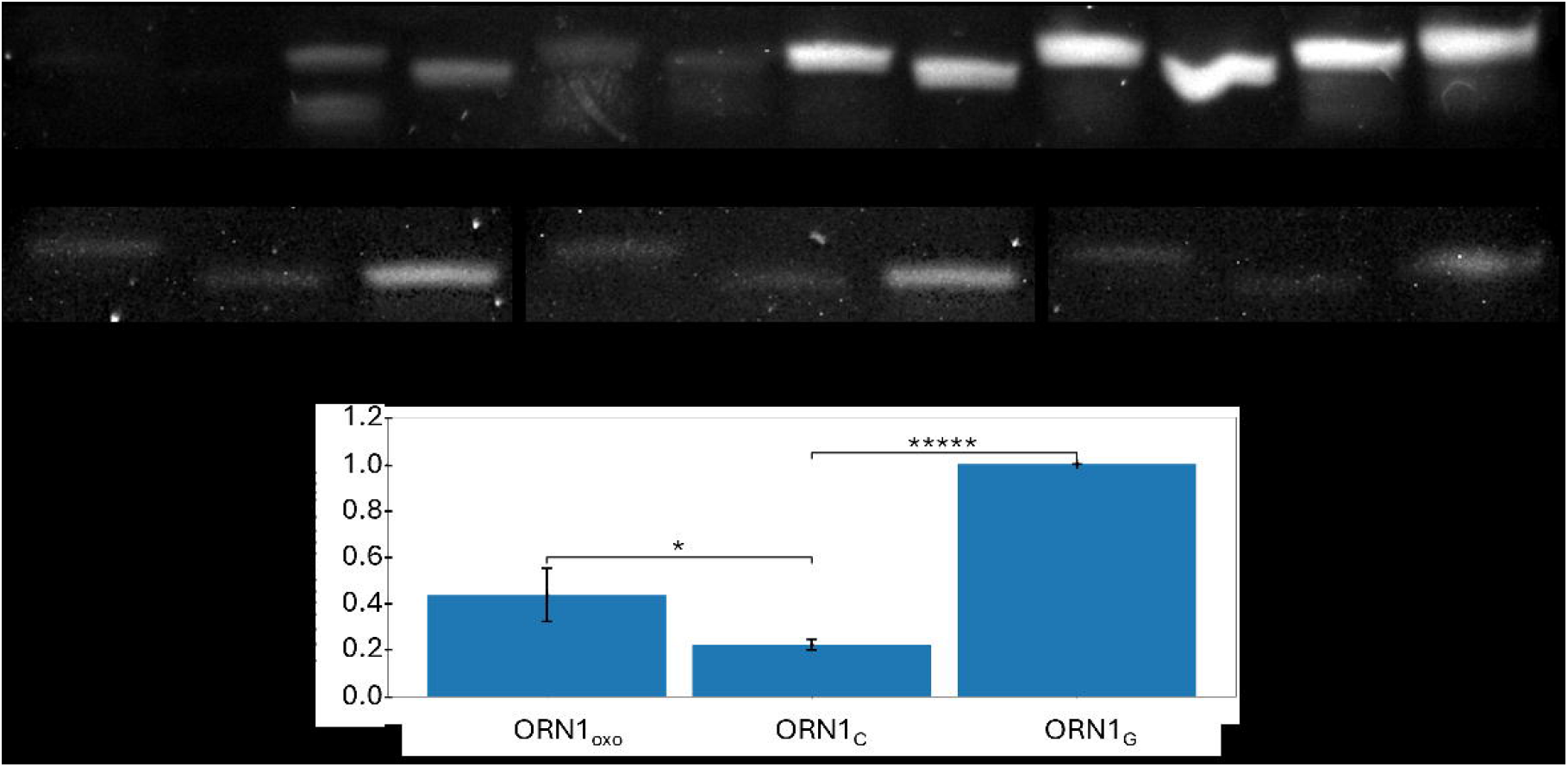
Reduction in SYBR Gold signal intensity for ribo-oligonucleotides with low content of unmodified guanosine. **A)** Imaged oligos 1-12 on the gel. It is apparent that oligos with fewer guanosines give much weaker signals, with no measurable signal visible for the oligo lacking any unmodified guanosine. The imaging method is described in the Methods section. **B)** Gel images (in triplicate) of three RNAs: ORN1_oxo_, ORN1_C_, ORN1_G_ bearing, in order: 8-oxo-G, C- or G-at indicated positions in the sequence (see **Tab.1**). 500 ng of oligonucleotide were used for each replicate. **C)** Quantification of band intensities in the gel image from B) using the ImageJ software. The intensities are normalized to the strongest signal, that of ORN1_G_. Error bars represent standard deviations, and the stars indicate cutoffs on the p-value from Student’s t-test: * p= 0.05; ***** p= 0.000005.

To test our observation, we selected the RNA from **Fig.1A (lane 1 and 2)** for which this effect was most pronounced (ORN1_oxo_ in **Table 1**) and procured oligonucleotides containing (1) only unmodified Gs (ORN1_G_, **Table1**); or (2) cytosines (Cs) in place of Gs (ORN1_C_, **Table 1**). Upon submitting equal amounts of all three oligonucleotides to electrophoresis on 14% urea PAGE (denaturing) gel and visualizing by SYBR Gold (**Fig. 1B**), we observed that the G-containing RNA oligonucleotide ORN1_G_ yielded approximately 5-fold higher signal intensity than the C-containing variant ORN1_C_ and approximately 2-fold higher than the RNA with oxidized Gs (ORN1_oxo_) **(Fig. 1C)**. This observation lent credence to our initial hypothesis that the disparity in signal intensity was driven by the unmodified guanosines.

**Table 1.**
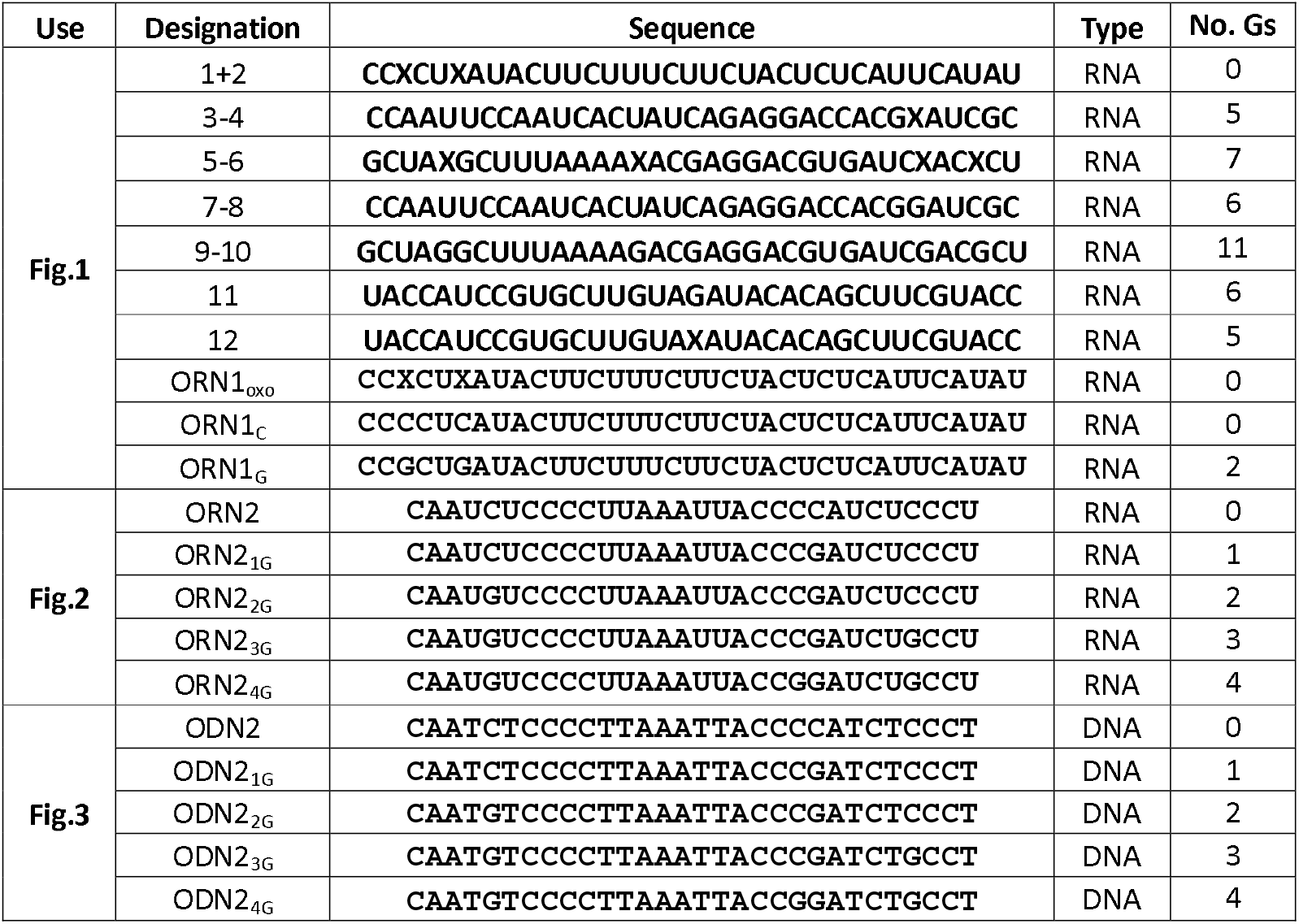
Sequences used in the experiments; X indicates 8-oxoguanosine (8-oxo-G).

Next, we sought to determine whether the SYBR Gold signal increases with the number of guanosines in the sequence, or whether the mere presence of guanosine is sufficient for the effect. To this end, we prepared five RNA oligonucleotides starting from the parent sequence ORN2 and replacing sequentially 1-4 cytidines by guanosines (**Table 1**). We repeated the experiments four times (**Fig. 2A)**, normalizing the measured signal intensities to that from the oligonucleotide containing four guanosines (ORN2_4G_) **(Fig. 2B)**. We found that signal intensity increased progressively with the guanosine content, the largest increase occurring upon the addition of the fourth guanosine (ORN2_3G_ = 0.28; ORN2_4G_ = 1.00). Overall, the signal intensity increased about 25-fold between ORN2 (no guanosines) and ORN2_4G_, the variant with four guanosines (ORN2 = 0.038; ORN2_4G_ =1.00). That the number of guanosines in a sequence influences SYBR Gold staining intensity further strengthens the hypothesis that guanosine is essential to dye binding and subsequent increase in fluorescence emission for single stranded RNA oligos.

**Fig. 2.**
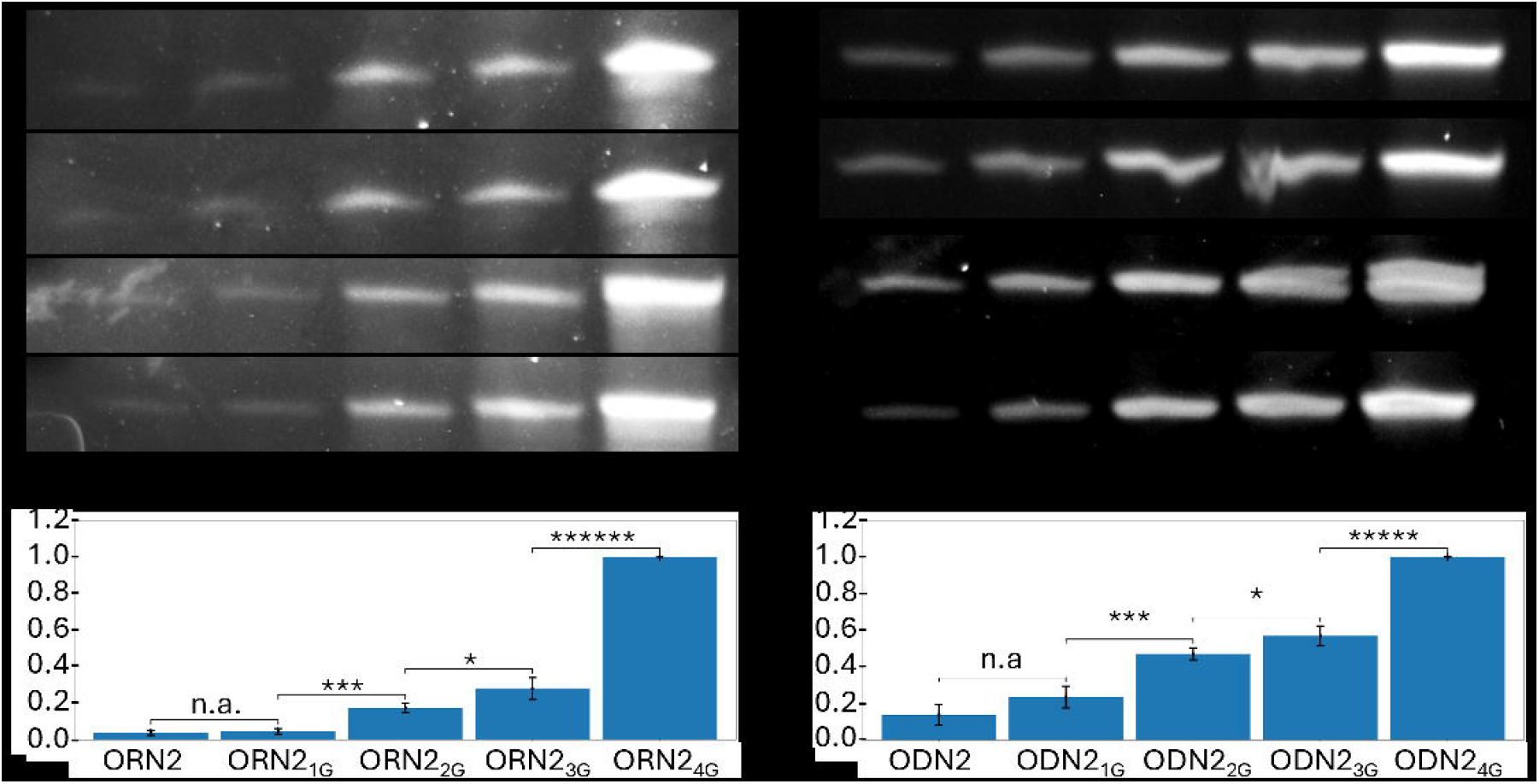
Signal intensity from SYBR Gold staining in ribo- and deoxyribo-oligonucleotides increases with guanine content. **A**) Denaturing gel images show the intensity of RNA oligonucleotide bands when stained with SYBR Gold in four replicates (rows). Oligonucleotides contain, from left to right, 0-4 guanosines (see **Tab. 1** for sequences). 500 ng of each oligo were used for each experiment. **B)** Quantification of band intensities in the gel image using the ImageJ software. The intensities are normalized to the signal of ORN2_4G_. Error bars represent standard deviations, and the stars indicate cutoffs on the p-value from Student’s t-test with n.a. indicating p>0.05, * p= 0.05 and ****** p= 0.0000005. **C)** Gel images show the intensity of deoxyoligonucleotide bands when stained with SYBR Gold (each row is a replicate experiment). Oligonucleotides contain, from left to right, 0-4 deoxyguanosines (see **Tab. 1** for sequences). 500 ng of each oligonucleotide were used. **D)** Quantification and representation of band intensities done in the same why as B. The intensities are normalized to the signal of ODN2_4G_.

To determine whether these observations extended to single-stranded DNA, we repeated the experiments using the same nucleotide sequence as for the RNA and denaturing gel **(Figs. 2C-D)**. The observations paralleled those made on RNA, with an even clearer correlation between fluorescence signal intensity and deoxyguanosine count. The relative intensity of the signal increased 7-fold from the parent DNA oligonucleotide with no deoxyguanosines (ODN2) to the DNA with four deoxyguanosines (ODN2_4G_) (ODN2=0.13; ODN2_4G_ =1) (**Fig. 2D**). Of note, the DNA sequence with no deoxyguanosines yielded a stronger signal than the corresponding RNA oligonucleotide.

Having shown that the guanosine content substantially impacts the signal intensity of SYBR Gold fluorescence in both RNA and single-stranded DNA contexts, we asked whether deoxyguanosine also drives the signal from double stranded DNA (dsDNA), where intercalation is thought to be the primary binding mechanism. Hence, we generated two dsDNA sequences: one composed exclusively of adenosine and thymidine, and the other containing all four deoxynucleotides (**Fig. 3A**). We observed no difference in the SYBR Gold fluorescence signal when staining equal amounts of these two dsDNAs **(Fig. 3A-B)**. This indicates that deoxyguanosine content is not a critical factor for the signal intensity of SYBR Gold in the context of helical structures.

**Fig. 3.**
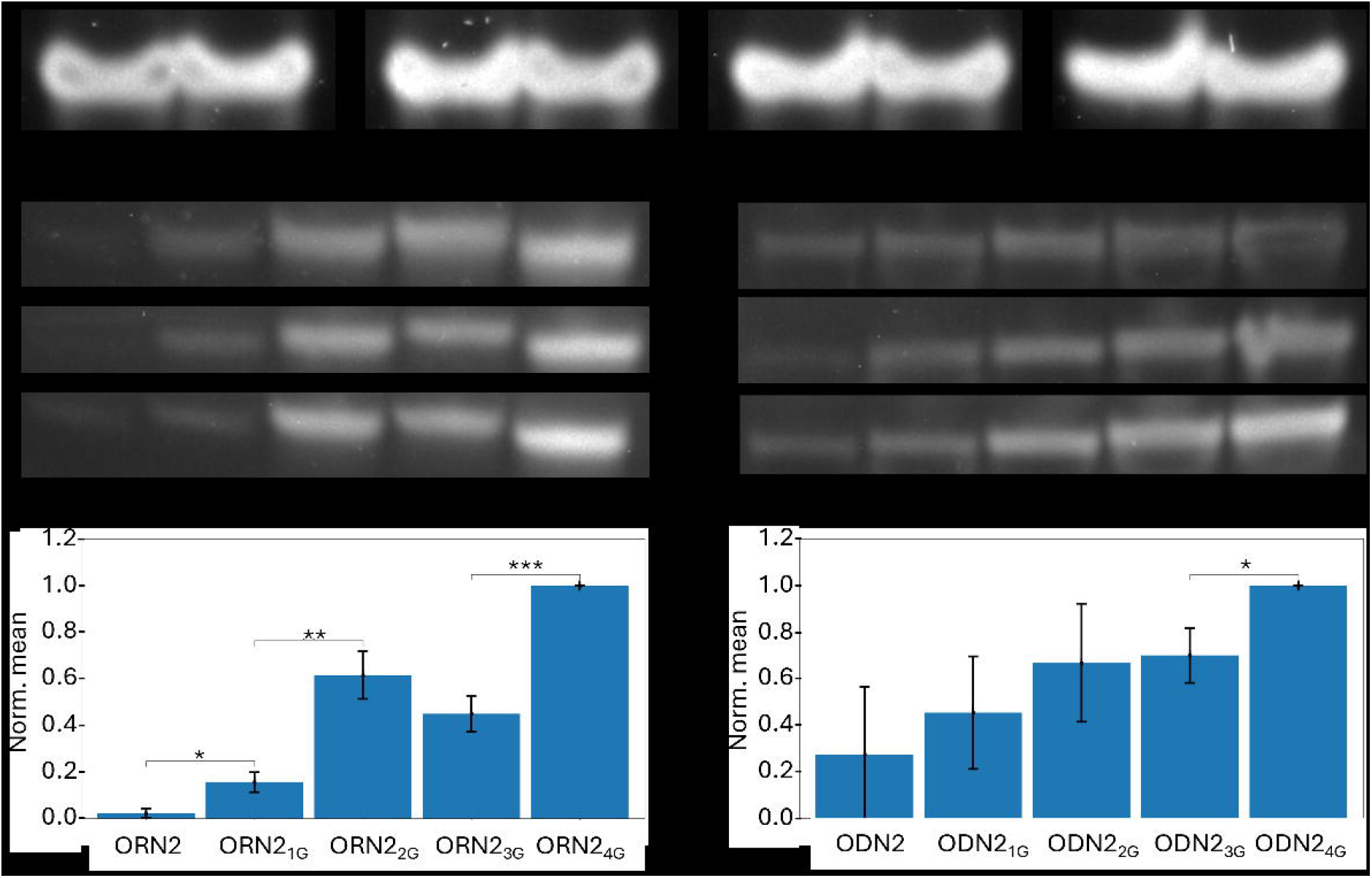
G-content dependency of SYBR Gold signal from single-stranded nucleic acids on native PAGE gels. **A)** Sequences of the dsDNA oligonucleotides used in the experiment. **B)** Four replicate (R1-R4) experiments using dsDNA probes containing only A/T on the left and A/T + C/G on the right. After running the samples on a 14% non-denaturing PAGE gel, no difference in signal staining intensity was observed. **C)** Three replicates of the RNA samples used in **Fig.2** run on a nondenaturing PAGE gel like the one described in B**). D)** Analysis of the data shown in **C)**. Error bars represent standard deviations, and the stars indicate cutoffs on the p-value from Student’s t-test with * indicating p= 0.05 and *** p= 0.0005. **E)** Three replicates of the DNA samples used in **Fig.2** run on a nondenaturing PAGE gel in parallel with the RNA samples. **F)** Analysis of the data shown in **E)**

One major difference between the experiments with dsDNA and ssDNA/RNA oligonucleotides are the Gel running conditions (native or denaturing respectively). To determine whether this factor can explain the some of the differences in fluorescence intensity, we performed native PAGE with RNA **(Fig.3C-D)** and ssDNA **(Fig. 3E-F)** oligonucleotides. The RNA oligos used in **Fig. 2** showed a similar trend of increasing signal as a function of guanosine abundance in these conditions, with a less monotonic increase of the signal with the number of Gs in the sequence **(Fig. 3C-D)**. The pattern was similar for the ssDNA oligonucleotides **(Fig. 3E-F)**, though the variability between replicates was clearly larger in the native PAGE compared to denaturing gels, despite identical experimental conditions, concentrations and timing for all replicates. Thus, while the gel conditions impact to some extent the results, native gels inducing a larger variability, the dependency of SYBR Gold signal intensity on the G content in single-stranded nucleic acids remains.

## Discussion

Taken together, our results indicated that the guanosine content strongly influences the fluorescence properties of SYBR Gold upon binding to single stranded nucleic acids in denaturing condition. In ssDNA the SYBR Gold signal increases almost linearly with the deoxyguanosine content (**Fig. 2C**), in line with the observed linear increase in fluorescence as a function of the number of SYBR Gold molecules bound to dsDNA^2^. While the signal intensities observed upon staining RNA did not appear to increase linearly, the overall increase in signal between the no-guanosine and four-guanosine sequence variants was higher. We speculate that the difference in baseline signal intensities between the guanosine-free RNA and deoxyguanosine-free DNA oligonucleotides may be due to an overall lower binding efficiency of SYBR Gold to RNA compared to DNA. Furthermore, we demonstrated that not only the absence of guanosine, but also its modification - in our case to 8-oxo-G (**Fig. 1B**) - can cause a loss of signal. This indicates a need for caution when using SYBR Gold in studies involving RNA modifications and denaturing gels.

In dsDNA, SYBR gold signal is independent of guanine content (**Fig. 2A – B**), likely being driven by intercalation, as described earlier^2^. While dsDNA is analyzed on native gels, ssDNA and RNA are analyzed on denaturing gels. To exclude this factor explaining the difference in signal intensity we repeated the experiments with ssDNA and RNA on native gels. The trend of increased signal with the G content remained, but the variability between replicates was higher, especially for ssDNA. Although the origin of this phenomenon remains to be determined, a previous study found that short (5-10nt) RNA oligos were unfolded in their native state while short ssDNA oligos with the same sequence were folded^3^. Thus, the higher propensity of ssDNA to form secondary structures and their stochastic dynamics may induce fluctuations in SYBR Gold interaction with the nucleic acid and, consequently, in signal intensity.

This study is limited to the analysis of SYBR Gold signals from a small number of short, synthetic oligonucleotides. Although any secondary structure in the sequences should not play a significant role when using a denaturing gel, studying a larger number of baseline sequences would provide a fuller picture of these previously unreported properties of SYBR Gold. We have not attempted to elucidate the specific mechanisms by which SYBR Gold binds to single-stranded RNA and DNA. However, the results presented here do suggest that guanosine contributes significantly to this mechanism. Our findings have important implications for the detection of single-stranded DNA and RNA, as they suggest that the intensity of SYBR Gold signals may depend more on guanosine content than on the overall abundance, making comparisons between oligonucleotides with different guanosine counts problematic.

In conclusion it appears that SYBR Gold may not be an optimal choice for visualizing ssDNA or RNA in denaturing gels particularly when modified guanines are present or when the guanosine content is low. Further using native gels will lead to much less consistent results especially for ssDNA samples.

## Author contributions

N.M. and M.Z. conceived the study. J.R. carried out the experiments. M.Zi. produces oligos. J.R. analyzed all data. J.R., N.M. and M.Z. wrote the manuscript with help from J.H. All authors have read and approved the manuscript.

## Acknowledgements

This work was supported in part by Jubiläumsstiftung von Swiss Life and by the NCCR RNA and disease. We acknowledge Dr. Adriano Biasini for his help and suggestions on the project.

